# Exogenous and endogenous HDAC inhibitor effects in Rubinstein-Taybi syndrome models

**DOI:** 10.1101/2020.03.31.015800

**Authors:** Elisabetta Di Fede, Emerenziana Ottaviano, Paolo Grazioli, Camilla Ceccarani, Chiara Parodi, Elisa Adele Colombo, Giulia Bassanini, Marco Severgnini, Donatella Milani, Elvira Verduci, Thomas Vaccari, Valentina Massa, Elisa Borghi, Cristina Gervasini

**Affiliations:** Department of Health Sciences, Università degli Studi di Milano, Milan, Italy; Institute of Biomedical Technologies, Italian National Research Council, Segrate, Milan, Italy; Fondazione IRCCS Ca’ Granda Ospedale Maggiore Policlinico, Milan, Italy; Department of Pediatrics Vittore Buzzi Children’s Hospital-University of Milan, Milan, Italy; Department of Biosciences, Università degli Studi di Milano, Milano, Italy; “Aldo Ravelli” Center for Neurotechnology and Experimental Brain Therapeutics, Università degli Studi di Milano, Milan, Italy

## Abstract

Rubinstein-Taybi syndrome (RSTS) is an autosomal dominant disorder with specific clinical signs and neurodevelopmental impairment. The two known proteins altered in the majority of RSTS patients are the histone acetylation regulators CBP and p300. For assessing possible ameliorative effects of exogenous and endogenous HDAC inhibitors (HDACi), we exploited *in vivo* and *in vitro* RSTS models. First, HDACi effects were tested on *Drosophila melanogaster*, showing molecular rescue. In the same model, we observed a shift in gut microbiota composition. We then studied HDACi effects in RSTS cell lines compared to healthy donor cells. We observed patients-specific molecular rescue of acetylation defects at subtoxic concentrations. Finally, we assessed commensal gut microbiota composition in a cohort of RSTS patients compared to healthy siblings. Intriguingly, we observed a significant depletion in butyrate-producing bacteria in RSTS patients. In conclusion, this study reports the possibility of modulating acetylation equilibrium by HDACi treatments and the importance of microbiota composition in a chromatinopathy.

## INTRODUCTION

Gene expression regulation is mediated by tightly balanced epigenetic mechanisms, such as histone modifications (as acetylation and methylation). Histone acetylation equilibrium on lysine residues is fundamental for several physiological processes, including correct embryonic development. Two classes of functionally antagonistic enzymes, acetyltransferases (HAT) and deacetylases (HDAC), are known to modulate this equilibrium (Grunstein, 1997). Histones hypoacetylation has been associated to alterations in synaptic plasticity, neuronal survival/regeneration, memory formation (Uchida and Shumyatsky, 2018); defects in epigenetic components acting on acetylation status cause several neurodevelopmental/malformation syndromes (Bjornsson, 2015). Among them, Rubinstein-Taybi syndrome (RSTS, OMIM #180849, #613684) is a rare (1:125,000) autosomal-dominant disease that occurs generally *de novo* (Rubinstein and Taybi, 1963), characterized by intellectual disability (Hennekam, 2006), postnatal growth deficiency with excessive weight gain in adolescence, distinctive dysmorphisms, skeletal abnormalities with a wide spectrum of multiple congenital anomalies (Hennekam, 2006).

RSTS is caused by pathogenic variants in one of two highly conserved genes: *CREBBP* (16p13.3) coding for cAMP response element binding protein (CREB) binding protein (also known as ‘CBP’) and *EP300*, coding for E1A-associated protein p300, mapping on chromosome 22q13.2. *CREBBP* is considered the “major gene” as found mutated in >50% RSTS patients while *EP300* gene mutations have been described in a minor fraction of patients (Fergelot et al., 2016).

CBP and p300 are ubiquitously expressed homologous proteins belonging to the lysine acetyl transferases (HAT) family (Valor et al., 2013), acting as co-factors of transcription, and required in multiple pathways of cell growth control, DNA repair, cell differentiation, and tumour suppression (Chan and La Thangue, 2001; Dutto et al., 2018; Kung et al., 2000; Oike et al., 1999; Tillhon et al., 2012; Yao et al., 1998). Their enzymatic activity enables the opening of chromatin leading to gene expression when acetylation targets histone tails (Chan and La Thangue, 2001; Weinert et al., 2018; Yao et al., 1998).

In recent years, a novel class of compounds, termed HDAC inhibitors (HDACi), have been used in different pathologies (Heerboth et al., 2014; Kazantsev and Thompson, 2008) for incrementing histone acetylation. Preliminary studies testing the efficiency of HDACi to revert acetylation defects in RSTS lymphoblastoid cell lines (LCLs) supported the hypothesis that RSTS is caused by acetylation imbalance (Lopez-Atalaya et al., 2012). Similarly, animal model studies supported the idea that chromatin alterations observed in RSTS could be reverted (Alarcón et al., 2004). In this complex and regulated equilibrium, another key component has been recently forwarded, in fact protein acetylation can also be modulated by the microbiota (i.e. commensal microbial community) (Simon et al., 2012). For example, short chain fatty acids (SCFAs, mainly acetate, propionate and butyrate) are the most abundant end-products deriving from anaerobic fermentation. SCFAs are pleiotropic microbial signals and, besides their key metabolic roles, they act as HDACi. Among them, endogenous butyrate is exclusively produced by commensal microorganisms and is the most potent HDACi among natural compounds (Stilling et al., 2016). Indeed, besides SCFAs production, an altered gut microbiota could participate in the typical RSTS growth trend (wasting deficit in infancy and excessive weight gain after puberty) and to comorbidities often associated to RSTS, such as gastrointestinal discomfort (Milani et al., 2015; Spena et al., 2015).

On these premises, in the present study we have exploited the experimental model *Drosophila melanogaster*, by using CBP mutant flies, to assess *in vivo* the effect of HDAC inhibition and for evaluating the genetic-determined microbiota characteristics in our insect model system. We have tested different HDACi molecules *in vitro* on lymphoblastoid cell lines (LCLs) derived from RSTS patients, to evaluate their effectiveness in modulating the previously assessed acetylation impairment by antagonizing the CBP/p300 defects (Lopez-Atalaya et al., 2012). Finally, microbiota-derived endogenous HDACi molecules have been evaluated by studying the gut microbial community in RSTS patients.

## RESULTS

### *HDACi exposure leads to acetylation increase in mutated* Drosophila *CBP homolog (*nej)

We exploited a RSTS *in vivo* model for assessing exogenous HDACi effects. Hence, we evaluated acetylation levels in *Drosophila* mutants for the CBP homolog *nejire* (*nej*) upon feeding with HDACi. To this end, heterozygous flies carrying a copy of the insertional mutant *nejP* (*nejP/+*) or yellow white (*yw*) control flies were reared on fly food supplemented with 1mM and 2.5mM VPA, or 10mM and 20mM NaB, or H_2_O as a vehicle. Proteins were extracted from 50 F1 female flies for each experimental condition. We performed Western blot analysis and quantifying levels of acetylated histone H3 and of β-Actin for normalizing purposes. As expected, extracts of *yw* flies show higher H3 acetylation compared to those of *nejP/+* animals supplemented with water (data not shown), while a different trend is present for extracts of flies fed with HDACi (Figure 1A and B). Interestingly, extracts of *nejP/+* flies show an increase in acetylated H3 when supplemented with 1mM VPA, 2.5mM VPA or 20mM NaB relative to *yw*.

**Figure 1.**
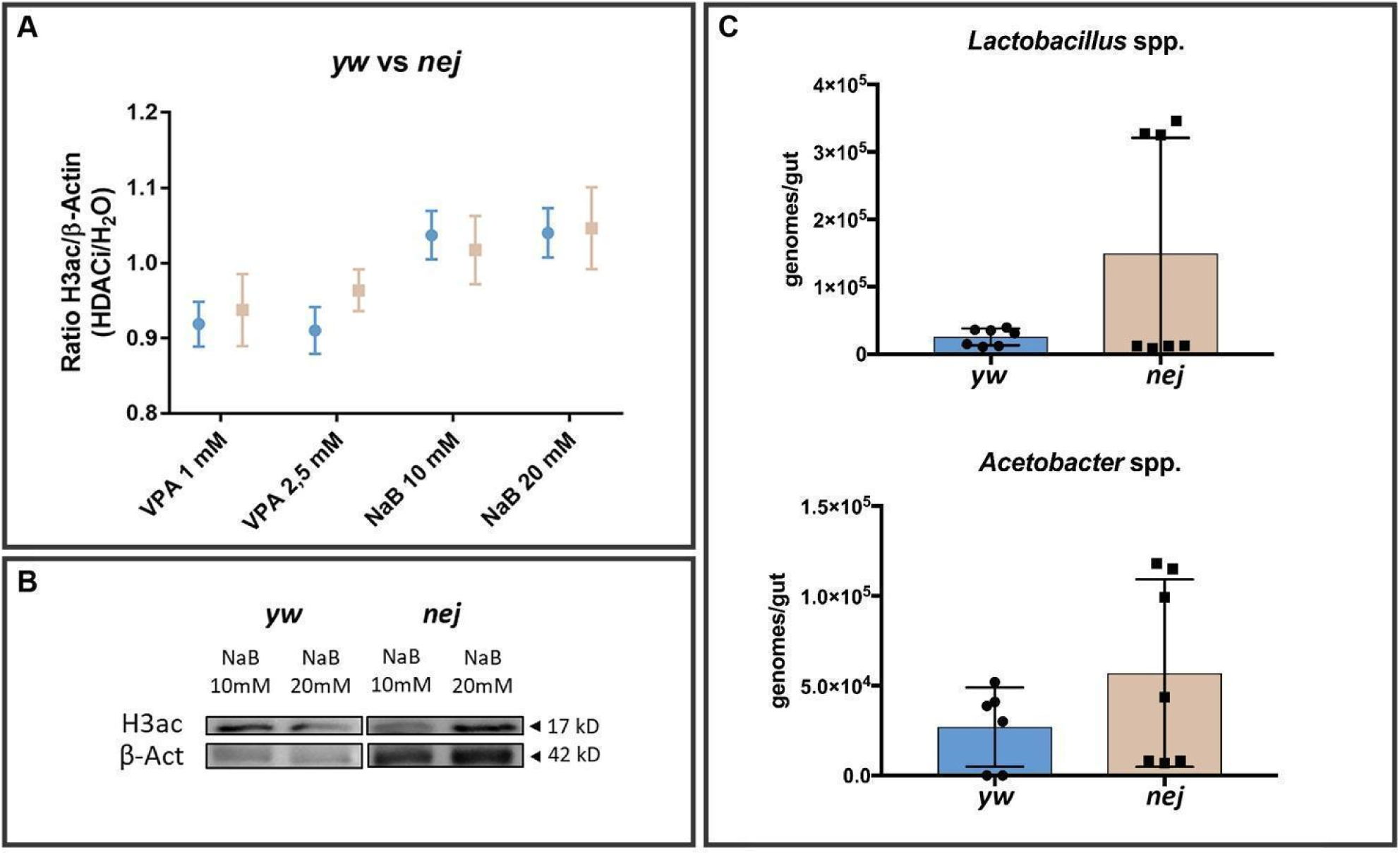
RSTS *Drosophila analysis of HDACi* molecular rescue and gut microbiota composition. H3 acetylation levels normalized on β-Actin, assessed with western blot analysis. A) Quantification of immunoblotting signals of H3 acetylation normalized and expressed as HDACi/H_2_O (Y-axis) in yellow white control flies (*yw*, in blue) and mutated Drosophila CBP homolog flies (*nej*, in pale brown), both fed with two different epigenetic treatments at two different doses (VPA 1mM, VPA 2,5mM, NaB 10mM and NaB 20mM, on X-axis); error bars represent means ± SEM and groups were compared using Student’s t-test as statistical method (significant value for p<0.05). B) Signals of *yw* and *nej* proteins immunoblotted with antibodies anti histone H3 (acetyl K9+K14+K18+K23+K27) and anti β-Actin after NaB 10mM and 20mM feeding. C) *Lactobacillus* spp. and *Acetobacter* spp. absolute quantification by realtime PCR in *yw* (blue) and *nej* (pale brown) flies. Abundances are expressed as genome copies/gut. Differences recapitulate those observed in RSTS patients (violin diagram for alpha-diversity in Supplementary file S7).

### *Mutations in the* Drosophila *CBP homolog (*nej*) increase inter-individual variation in the gut microbial community*

As *Drosophila* laboratory-reared strains display a very simplified gut bacterial community (Douglas, 2018; Wong et al., 2013), the two most represented genera, *Lactobacillus* spp. and *Acetobacter* spp., were quantified by Real-time PCR (Figure 1C). We did not observe statistically significant differences in the abundance of these taxa between *nejP/+* and *yw* flies. Importantly, we noticed an increased individual-to-individual variability in mutant flies. The coefficients of variation for *Lactobacillus* spp were 48.8% and 115.4% for *yw* and *nejP/+*, respectively; for *Acetobacter* spp. 81.2% and 91.3%, respectively.

### *HDACi exposure counteracts* acetylation imbalance in RSTS lymphoblastoid cell lines (LCLs)

Having confirmed the possible ameliorative effects in our *in vivo* model, we then sought to study HDAC inhibition in patients-derived cells. We exposed LCLs derived from eight RSTS patients with *CREBBP* (n.4) or *EP300* (n.4) confirmed mutations (Supplementary file S1) and seven healthy donors (HD) to four different HDAC inhibitors (HDACi): Trichostatin A (TSA), Suberoylanilide hydroxamic acid (SAHA), Valproic acid (VPA) and Sodium Butyrate (NaB) (Supplementary file S2). We analysed by AlphaLISA^®^ assay the acetylation levels of lysine 27 of histone H3 (H3K27ac) in LCLs upon three different conditions: HDACi treatments, exposure to vehicle (DMSO or H_2_O) and untreated cells (Figure 2).

**Figure 2.**
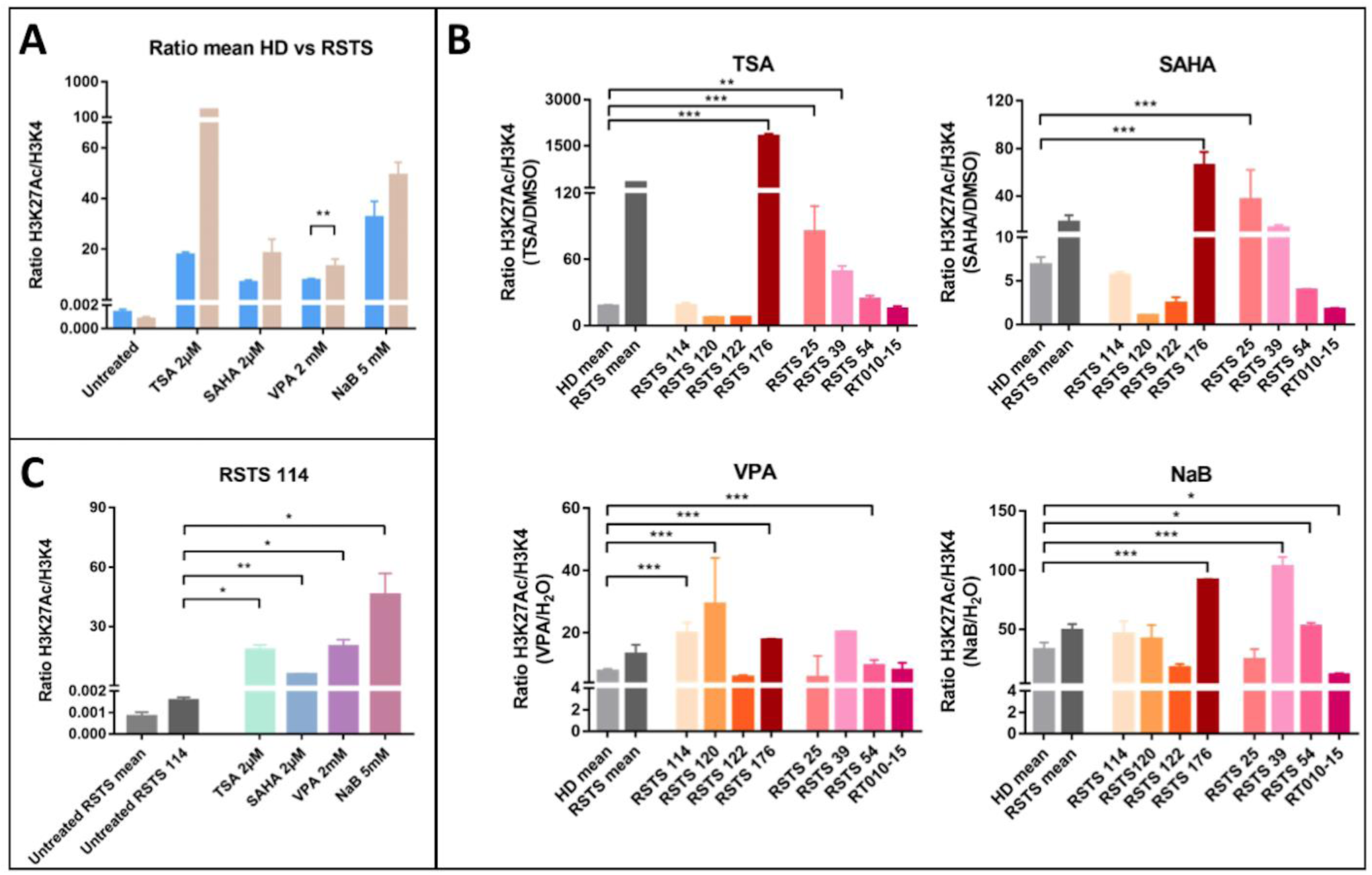
Histone acetylation on RSTS LCLs upon HDAC inhibitors exposure. H3K27 acetylation levels normalized on H3K4 unmodified, assessed by AlphaLISA^®^; levels of acetylation upon HDACi are expressed as ratio between the treatment and respective vehicle; on Y-axis H3K27 acetylation levels normalized and expressed as HDACi/vehicle, on X-axis lists of epigenetic treatments or untreated/treated single LCL or LCLs means A) Means of values of H3K27 acetylation in healthy donors (HD, in blue) and patients LCLs (RSTS, in pale brown) untreated and exposed to four different HDACi (TSA 2μM, SAHA 2μM, VPA 2mM and NaB 5mM). B) H3K27 acetylation in eight RSTS LCLs (*CREBBP* LCLs in shades of red, *EP300* LCLs in shades of pink) after exposure with the four different HDACi, compared to treated HD and RSTS means. C) Insight on single-patients’ response (RSTS 114) to the four compounds compared to untreated RSTS means and RSTS 114. Groups were compared using Student’s *t-test* as statistical method (*p<0.05; **p<0.01; ***p<0.001).

All the compounds succeeded in boosting histone acetylation in RSTS LCLs compared to HD LCLs, with VPA exposure resulting highly significant (p<0.01). This increment was particularly manifest in patient-derived LCLs compared to untreated samples (Figure 2A).

We also observed that HDACi compounds induced a variable acetylation response, in a patient-specific manner when compared to treated HD LCLs (Figure 2B). As shown in figure 2B, treatment with TSA 2μM boosted significantly acetylation levels in LCLs RSTS 176 (p<0.001), RSTS 25 (p<0.001) and RSTS 39 (p<0.01), while SAHA 2μM showed highly significant effect on RSTS 176 and RSTS 25 (p<0.001). VPA 2mM treatment particularly increased H3K27ac of RSTS 114, RSTS 120, RSTS 176 and RSTS 54 (p<0.001), while exposure to NaB 5mM significantly affected acetylation of RSTS 176 and RSTS 39 (p<0.001), RSTS 54 and RT010-15 (p<0.05).

To note, when analysing specific RSTS patient-derived LCLs response to HDACi compared to the relative untreated conditions, we observed that at least one HDACi significantly boosted acetylation and that RSTS-LCLs response varied among different drug treatments (Figure 2C and Supplementary file S3).

### HDACi exposure does not affect cell-cycle in RSTS-derived LCLs

In order to investigate HDACi effect on cell-cycle regulation, cell proliferation and cell death were assessed upon HDACi exposure with, respectively, Ki67 and Tunel assays. For both assays, we did not observe a significant correlation with H3K27 acetylation (Figure 3).

**Figure 3.**
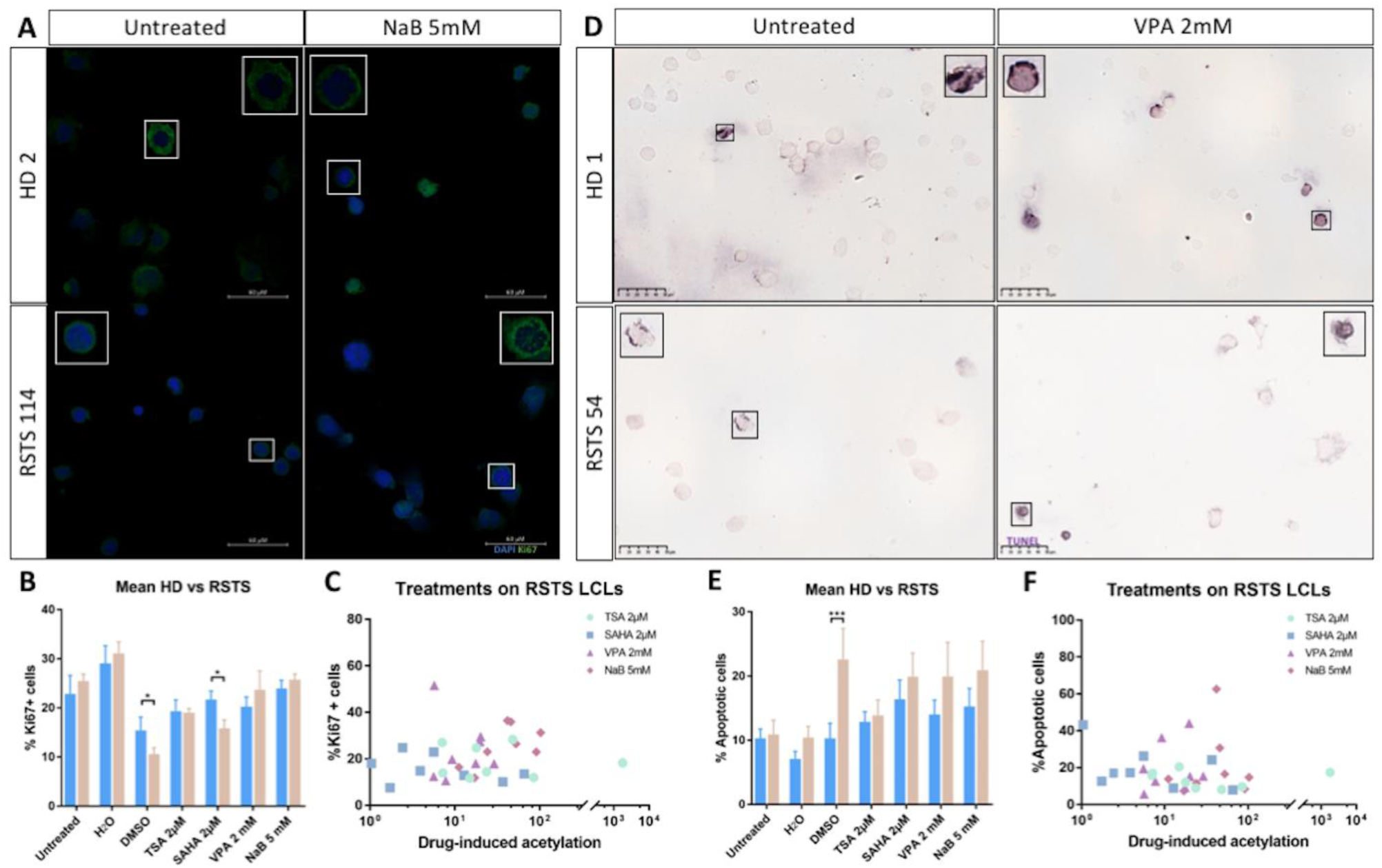
HDAC inhibitors cytotoxicity analysis on RSTS LCLs. Cell proliferation (A-C) and cell death (D-F) in RSTS LCLs compared to HD LCLs. A) Confocal 60x images showing an example of Ki67 assay performed on HD LCL (HD2) and RSTS LCL (RSTS 114) untreated and after exposure to NaB 5mM; nuclei are marked with DAPI (blue) and proliferative cells with Ki67 (green signal); Insets show 100x cell magnification. B) Cell proliferation rate of Ki67 positive cells (% Ki67+ cells, on Y-axis) of RSTS LCLs compared to HD LCLs upon HDACi exposure (TSA 2μM, SAHA 2μM, VPA 2mM and NaB 5mM), treatment with vehicles (H_2_O and DMSO) and untreated condition (X-axis). C) Correlation overview between cell proliferation rate (% Ki67+ cells, on Y-axis) and drug-induced acetylation (X-axis) in RSTS LCLs exposed to different HDACi (TSA 2μM, SAHA 2μM, VPA 2mM and NaB 5mM). D) Brightfield 40x acquisitions showing an example of TUNEL assay performed on HD LCL (HD1) and RSTS LCL (RSTS 54) untreated and after exposure to VPA 2mM, with apoptotic cells appearing deep purple (TUNEL positive); Insets show 80x cell magnification. B) Cell death rate of TUNEL positive cells (% Apoptotic cells, on Y-axis) of RSTS LCLs compared to HD LCLs upon HDACi exposure (TSA 2μM, SAHA 2μM, VPA 2mM and NaB 5mM), treatment with vehicles (H_2_O and DMSO) and untreated condition (X-axis). C) Correlation overview between cell death rate (% Apoptotic cells, on Y-axis) and drug-induced acetylation (X-axis) in RSTS LCLs exposed to different HDACi (TSA 2μM, SAHA 2μM, VPA 2mM and NaB 5mM). Cell proliferation and cell death rate groups were compared using Student’s *t-test* as statistical method (*p<0.05; **p<0.01; ***p<0.001).

Ki67 assay revealed no significant differences in proliferation rate between RSTS and HD LCLs except for exposure to vehicle DMSO and SAHA 2μM (p<0.05) (Figure 3A-B), however variable proliferation was observed in response to HDACi treatments among different RSTS LCLs (Supplementary file S4). We found no correlation between cell proliferation and drug-induced acetylation (Pearson correlation coefficient <0.3; Figure 3C). In details, treatments with TSA 2μM and VPA 2mM showed a very weak negative correlation (r=-0.03 and r=-0.11 respectively), SAHA 2μM a weak negative correlation (r=-0.3), while NaB 5mM a moderate positive correlation (r=0.45) (Supplementary file S4).

TUNEL assay (Figure 3D-F) showed significant differences in cell death for patients LCLs exposed to DMSO compared to HD LCLs (Figure 3E), as expected (Supplementary file S4). Importantly, no significant correlation was observed between apoptosis rate and HDACi-induced acetylation (Figure 3F): TSA 2μM and VPA 2mM showed, respectively, a weak and a very weak positive correlation (r=0.3 and r=0.11), while SAHA 2μM and NaB 5mM shared a very weak negative correlation (r=-0.038 and r=-0.06 respectively) (Supplementary file S5).

### *RSTS patients are depleted in the major butyrate-producer* Faecalibacterium *spp*

Having observed a difference in microbiota composition in our *in vivo* model, we enrolled 23 RSTS subjects (mean age 10.2 ± 6.4 years; 12 females) and 16 healthy siblings (healthy donors, HD), mean age 12.7 ± 7.2 years; 6 females), as a control group to minimize environmental factors having a well-recognized role on gut microbiota. The dietary survey revealed no differences for all macronutrients but energy intake, lower in RSTS (p=0.0054). Nutritional parameters are detailed in the relative supplementary table (Supplementary file S6).

Microbiota profiling was performed by 16S rRNA gene-targeted sequencing. After quality filtering processes, we obtained a mean count of 90,759 reads per sample. The alpha-diversity analysis of the gut microbiota showed no significant differences between RSTS and HD faecal samples in term of richness (Observed species: p=0.255; PD-whole tree: p=0.279 - Figure 4A) and richness and evenness (Chao1: p=0.151; Shannon: p=0.287-Supplementary file S7). Beta-diversity analysis, instead, showed that RSTS faecal microbiota differed significantly from that of healthy group according to both unweighted (p=0.013) and weighted (p=0.022) Unifrac distances (beta-diversity, Figure 4B).

**Figure 4.**
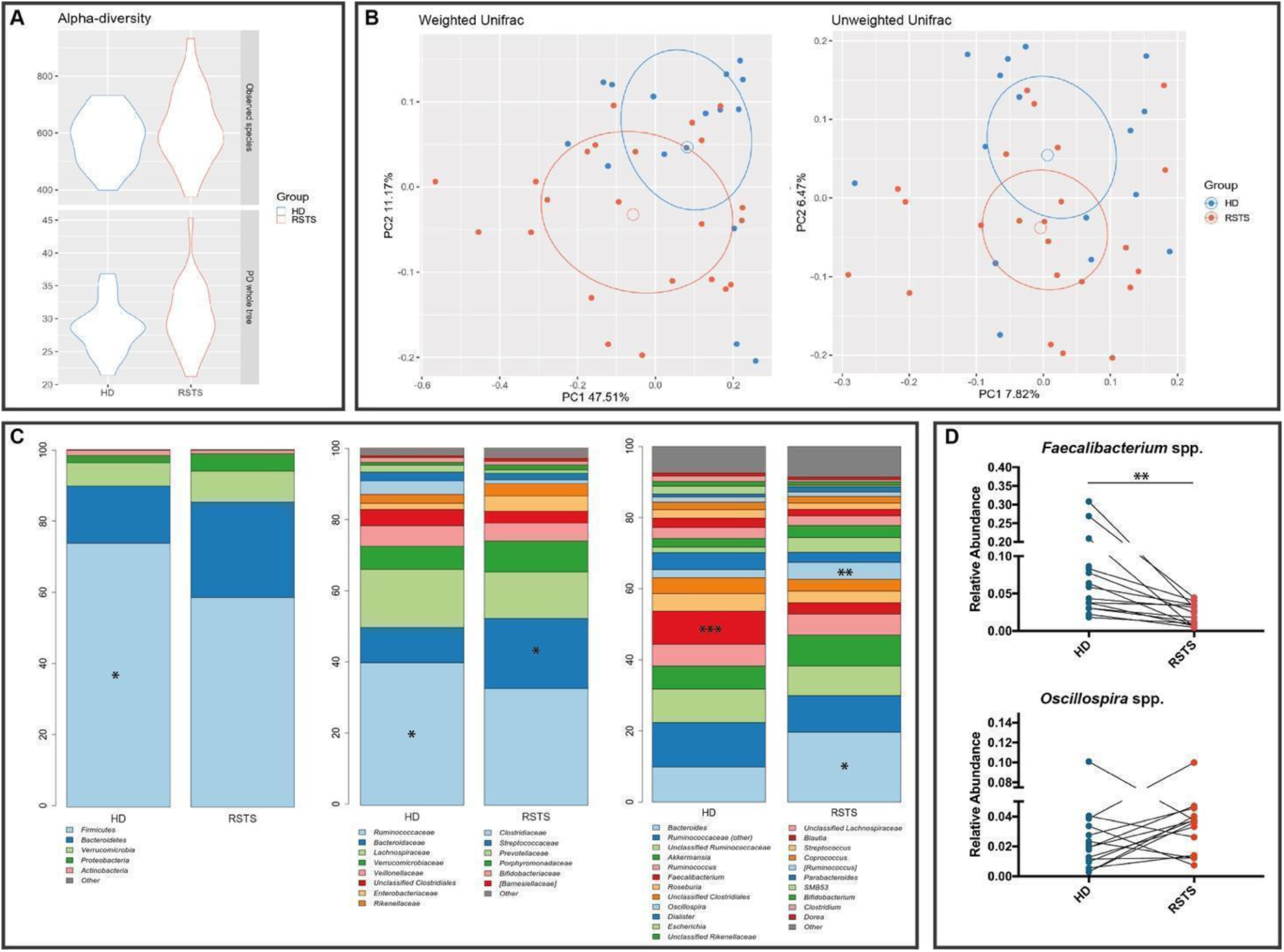
RSTS gut microbiota analysis. Gut microbial community was characterized by 16S rRNA gene sequencing. A) alpha-diversity. The violin plot shows biodiversity values for observed_specied and Faith’s phylogenetic metrics. No statistically relevant differences were seen. B) Principal Coordinate Analysis (PCoA) according to weighted and unweighted Unifrac distances. Microbial communities are statistically different (*adonis* test: unweighted *p*=0.019; weighted *p*=0.023). First and second principal coordinates are shown in the plot for both distances. C) Bacterial composition of HD and RSTS groups. Relative taxonomic abundances are shown at phylum, family and genus phylogenetic levels. All bacterial taxa present at <1% relative abundance were grouped into the “Other” classification. ***: p<0.005; **: p<0.01; *: p<0.05. D) *Faecalibacterium* spp and *Oscillospira* spp relative abundances (both significantly different between RSTS and HD) were compared within matched family members (patient/sibling, n=16). Whereas for *Oscillospira* we did not observe a common pattern, *Faecalibacterium* spp. was significantly reduced in RSTS (p=0.0021, Wilcoxon signed rank test).

The overall composition of the intestinal microbiota (Figure 4C; Supplementary file S8), showed a decreased relative abundance of the Firmicutes phylum (58.5% in RSTS *vs* 73.4% in HD, p=0.019), of the *Ruminococcaceae* family (32.2% vs 41.9%, p=0.049) and of *Faecalibacterium* spp. (3.3% *vs* 9.8% in HD, p=0.001) in RSTS subjects. On the other hand, RSTS samples showed an enrichment in the *Bacteroidaceae* family and in *Bacteroides* spp. (21.1% *vs* 10.3%, p=0.021) as well as in *Oscillospira* spp. (5.1% *vs* 2.4% in HD, p=0.007). Matched-pair analysis (Wilcoxon signed-rank test), performed on RSTS/sibling pairs showed a significant and environment-independent decrease (p=0.0021) in *Faecalibacterium* spp. (Figure 4D).

Short-chain fatty acid measurements showed a slight decrease in faecal butyrate content (4.13 ± 1.40 vs 5.14 ± 1.79 mg/g faeces, p=0.0741, Mann-Whitney test), whereas acetate, propionate, and branched-chain fatty acids (iso-butyrate and iso-valerate) concentrations were similar (p=0.194, p=0.874, p=0.786, and p=0.467, respectively).

## DISCUSSION

The current therapeutic approach for RSTS patients is directed towards alleviating clinical symptoms and preventing possible known comorbidities. For example, common interventions for these patients are behavioural support and surgical procedures for the correction of orthopaedic or cardiac malformations. In this context, exploring known drugs effects in preclinical studies is the fundamental step for envisaging therapeutic avenues (Cobos et al., 2019; Soragni and Gottesfeld, 2016). Hence, the present work explored the effects of exogenous and endogenous molecules in RSTS experimental models, that should rescue the defective enzymatic activity underlying the condition.

First, to study the effects of exogenous HDAC inhibition *in vivo*, we established and exploited *Drosophila melanogaster* model. *nej* is the unique fly homolog of the highly conserved CBP and p300 paralogs. Studies with *nej* flies underline the role of CBP during embryogenesis and as coactivators of critical transcription factors for developmental patterning (Akimaru et al., 1997; Tanaka et al., 1997). In our system, rearing flies with food supplemented with selected HDACi showed partial rescue of hypoacetylation in *nejP/+* animals, confirming the acetylation increase in an *in vivo* insect model. Finally, having exploited the *in vivo* model, we decided to analyse the microbiota differences in CBP insects compared to wild-type animals. Lack of an anoxic compartment in the *Drosophila* gut shapes a microaerophilic microbiota constituted, in laboratory strains, by few genera (Capo et al., 2019). The most abundant taxa are Firmicutes, mainly *Lactobacillus* spp., and alpha-Proteobacteria, mainly *Acetobacteraceae*. Despite evolutionary divergence with the human microbiota, recent studies showed that an altered relative abundance of these genera can result in gut homeostasis disturbance (Fast et al., 2018), growth delay (Shin et al., 2011), and behavioural changes (Sharon et al., 2010). The relative simplicity of the *Drosophila* bacterial microbiota and the availability of *Drosophila* mutants for a variety of human diseases allow also studying the influence of host factors on microbiota (Broderick and Lemaitre, 2012). Indeed, our results showed that, accounting for the species-specificity, the differences observed in RSTS patients compared to healthy siblings are recapitulated in RSTS flies compared to wild-type animals, strengthening our findings on gene-microbiota intimate interactions. Using this modelling system, we confirmed the possible ameliorative effects of acetylation boosting and we unravelled a microbiota composition difference probably caused by genetic background. For this, we decided to expand the study using patients derived cells and recruiting RSTS patients for microbiota sampling. A number of HDACi currently used for other purposes (José-Enériz et al., 2019; Lipska et al., 2020; Spartalis et al., 2019; Tarnowski et al., 2019) have been thoroughly evaluated in cell lines derived from patients compared to healthy donors. Upon HDACi exposure, RSTS cell lines showed a general boost in histone acetylation levels. Such increment was significant in a patient-specific manner. Indeed, each line, derived from different RSTS patients with discrete pathogenic variants responded differently to the tested compound. This data provides evidence of HDACi capability of restoring acetylation levels in an *in vitro* model of RSTS, strongly pointing to tailored future perspective in accordance with the idea of personalized medicine. We also evaluated the cytotoxicity of selected concentrations, given that HDACi are also used in oncological trials for their ability to induce selected and dose-dependent apoptosis (De Schutter and Nuyts, 2009; Yuan et al., 2019). Importantly, we did not observe neither an increase in cell death nor a decrease in cell proliferation, indicating that HDACi can boost acetylation at sub-toxic concentration in our *in vitro* modelling system.

For ascertaining if endogenous HDACi producing commensal bacteria could play a role in this interlinked equilibrium, we designed a gut microbiota study in patients. Given that there is no information about gut microbiota composition and metabolite production in RSTS subjects, we sequenced the V3–V4 hypervariable regions of the 16S rRNA gene and measured the main microbial metabolites -SCFAs-. This step aimed also at elucidating whether a distinct microbial signature could participate in the syndrome comorbidity insurgence. To rule out a direct effect of dietary habits or environment in microbiota alteration, healthy siblings were enrolled as control group. Despite the fact that we recorded no differences in the nutritional parameters, with the exception of the daily energy intake, the gut microbial community of RSTS was significantly different compared with healthy siblings. Both groups showed similar biodiversity, but displayed a distinct gut microbial ecosystem. The most relevant shift involved the butyrate-producer genus *Faecalibacterium* (Louis and Flint, 2009), strongly depleted in the RSTS group. This data is extremely relevant, considering that butyrate acts as endogenous HDACi (Davie, 2003) and further study should be aimed to elucidate this possible genetic-microbial additive negative effect. A recent study reported the effects of a ketogenic diet in a mouse model of Kabuki syndrome (Benjamin et al., 2017). Kabuki syndrome (OMIM# 147920, # 300867) is a rare genetic syndrome that overlaps Rubinstein-Taybi syndrome both at the clinical and genetic levels. Patients share many clinical signs and, at the molecular level, both syndromes are due to defects in the epigenetic machinery, mainly involving the histone modification level. In the Kabuki mouse model, ketogenic diet induces the endogenous production of deacetylase inhibitors that normalizes the overall state of histone modification and determines an improvement of clinical conditions (Benjamin et al., 2017). Ketogenic diet, having an unbalanced macronutrient composition, highly impacts on both microbiota composition and metabolism (Lindefeldt et al., 2019). In summary, we report that exogenous HDACi show molecular rescue of hypoacetylation *in vivo* and *in vitro* and that microbiota composition is altered in RSTS models compared to the relative controls, especially in HDACi producing bacteria. To note, carbohydrates and in particular fermentable dietary fibres, the most important substrates for short-chain fatty acid production (Bach Knudsen et al., 2018), were very similar in the two groups, thus not accounting for neither the reduction in the relative abundance of *Faecalibacterium* genus nor for the lower butyrate faecal concentration. Nutritional recommendations for RSTS comorbidity management are currently lacking as no studies focused on this aspect. Our findings might represent a starting point for further evaluation of specific nutritional regimen, which could also shed light on the basis for the observed differences.

Our overall results strongly support further studies in HDACi drug repurposing considering that HDACi are widely used and have ascertained toxicodynamic and toxicokinetic. In addition, several authors suggest that brain functions and behaviour are influenced, through a bottom-up modulation, by gut microbiota (Bercik et al., 2011; Heijtz et al., 2011), thus future effort will also point to enhancement of endogenous HDACi production.

## MATERIALS AND METHODS

### Drosophila melanogaster *stocks and HDACi feeding*

*Drosophila yw* strain was used as control, while the *nej*^*P*^ mutant strain was provided by the Bloomington Stock Center: (stock #3728). Flies were maintained and raised into vials containing standard food medium composed of yeast, cornmeal, molasses, agar, propionic acid, tegosept and water. All the strains were kept at 25°C. To prepare food with HDACi, stock solutions of VPA (1 mM and 2.5 mM) and NaB (10 mM and 20 mM) solutions were diluted 1:10 in the food before solidification but under 65°C to prevent heat damage of the compounds.

### Drosophila *protein extraction and western blotting*

50 female flies reared on supplemented food or control per sample were frozen at −80°C for 10 min, then RIPA buffer with proteases inhibitor (Complete, MINI, EDTA-Free - Roche #04693159001) was added in the tubes. Flies were homogenized with a sterile pestle and the tubes were then centrifuged at 14,000 rpm for 20 min at 4°C. Supernatants containing proteins were collected and stored at −80°C.

Proteins extracted from *Drosophila* were quantified with BCA Protein Assay kit (Biorad, CA, US), 20μg per sample were prepared with 4x Laemmli Sample Buffer added with β-mercaptoethanol (#1610747 and #1610710, Biorad), boiled for 10 min at 100°C, loaded and separated on 4-15% precast polyacrylamide gel (#4561083, Biorad). The run was carried out with a constant voltage of 100V into a running buffer composed by 25mM Tris, 192mM glycine and 0,1% SDS. Transfer was performed with 100V constant for 2h at 4°C into a buffer composed by 25mM Tris, 192mM glycine and 20% methanol. After gel transfer to PVDF membrane (#1620177, Biorad), blocking of aspecific sites was carried out using 5% milk in TBS-T (20mM Tris, 150mM NaCl, 0,1% Tween, pH 7.5) for 1h at RT. Primary antibodies anti histone H3 (acetyl K9+K14+K18+K23+K27) (ab47915, abcam) and anti β-Actin (A2228, Sigma-Aldrich) were diluted in 5% milk in TBS-T (1:1,000 and 1:2,000 respectively) and incubated at 4 °C overnight. After three TBS-T washes, membrane was incubated with HRP-conjugated secondary antibodies anti-Rabbit and anti-Mouse (diluted 1:2,000 in TBS-T) for 1h at RT. The membrane was washes in TBS-T three times and incubated 5 min with Clarity Western ECL Substrate (#1705061, Biorad) in order to detect proteins immunoblotted signals at Alliance LD4 (UVItec Cambridge, UK) which were analysed with UVI-1D software (UVItec Cambridge, UK).

### Drosophila melanogaster *DNA extraction and bacteria quantification by real-time PCR*

Genomic DNA was extracted using the QIAamp DNA Microbiome Kit (QIAGEN, Hilden, Germany), according to manufacturer’s recommendations. Briefly, three dissected guts from *Drosophila* adults per experimental group were pooled. Experiments were run in triplicates. After a washing step with phosphate buffered saline (PBS), a depletion of host cells DNA step was performed by adding lysis buffer and benzonase to samples. Bacterial cells lysis was then achieved using Pathogen Lysis Tubes (containing beads) and a lysis buffer in the TissueLyser LT instrument (QIAGEN). Lysates were transferred to QIAamp UCP Mini Columns and bound DNA was eluted in 50 μL of elution buffer. Absolute quantification by real-time PCR was performed using the following control strains: *Lactobacillus reuteri* and *Acetobacter pomorum* from Clinical Microbiology Laboratory of the Department of Health Sciences, Università degli Studi di Milano. Total microbial DNA was extracted using Prepman Ultra (Applied Biosystems, Foster City, CA). qPCR was carried out using the StepOne Plus instrument (Life technologies) and SYBR Premix Ex Taq II (Tli RNaseH Plus, Takara Bio). The specific 16S rRNA primers and qPCR conditions used for *Lactobacillus* spp. are reported by Borgo et al. (Borgo et al., 2017), and for *Acetobacter* by Ryu et al. (Ryu et al., 2008). Standard curve was carried out for each qPCR run using five serial dilutions of control DNA. Samples and standards were run in triplicate. The genome copy number was determined using the formula:

Number of copies=(ng*Avogadro constant)/(genome length*dsDNA average molecular weight)

Genome length was determined by the Ribosomal RNA Database (https://rrndb.umms.med.umich.edu). For lactobacilli, whose genome ranges from 1.8Mbp to 3.3 Mbp, the mean value was applied.

### Cell Cultures

Lymphoblastoid cell lines (LCLs) from eight different RSTS patients (four carrying *CREBBP* mutations and four carrying *EP300* mutations, listed in supplementary file 1) and seven healthy donors were obtained in collaboration with the Gaslini Genetic Bank service (Telethon Network of Genetic Biobanks). Cells were maintained in RPMI 1640 culture medium supplemented with L-glutamine (Euroclone, IT), 20% foetal bovine serum (Euroclone, IT) and penicillin/streptomycin (Euroclone, IT), and cultured in an incubator with 5% CO_2_ at 37°C.

LCLs were exposed to four different HDAC inhibitors: Trichostatin A (TSA) (sc-3511, Santa Cruz Biotechnology), Suberoylanilide hydroxamic acid (SAHA) (MK0683, Selleckchem), Valproic acid (VPA) (P4543, Sigma Aldrich) and Sodium Butyrate (NaB) (B5887, Sigma-Aldrich). We tested three different concentrations for each HDACi (Supplementary file 2) and selected the maximum dose and timing of exposure ensuring acceptable LCLs survival (data not shown). Cells were incubated with vehicles (H_2_O or DMSO) at maximum time (24h), TSA 2μM for 2h, SAHA 2μM for 24h, VPA 2mM for 24h or NaB 5mM for 24h as suggested from the literature (Supplementary file 2) (Chang et al., 2018; Chriett et al., 2019; Freese et al., 2019; Gottlicher M et al., 2001; Schölz et al., 2015; Tarasenko et al., 2018).

### AlphaLISA^®^ assay

After treatments, lymphoblastoid cellular pellets were obtained by centrifugation and frozen at −80°C. An amount of ~10,000 cells/well resuspended in 60 μl of culture media was used in order to perform AlphaLISA^®^ assay (PerkinElmer, MA, US) according to manufacturer’s protocol. Briefly, cells were incubated 15 min with Cell-Histone Lysis buffer and 10 min with Cell-Histone Extraction buffer; 30 μl of lysates were incubated with 10 μl of Acceptor mix 1h at RT and then 10 μl of Donor mix was added overnight at RT. Replicates were tested with both AlphaLISA Acetylated-Histone H3 Lysine 27 (H3K27ac) Cellular Detection Kit (AL720, PerkinElmer) and AlphaLISA unmodified Histone H3 Lysine 4 (H3K4) Cellular Detection Kit (AL719, PerkinElmer) for normalization. PerkinElmer EnSight(tm) plate reader was used for the detection of the chemiluminescent signal.

### Ki67 and TUNEL assay

After treatments, at least 1.5×10^4^ LCLs were seeded in duplicate on SuperFrost Plus slides (Thermofisher Scientific, MA, US) through 5 min of cytospin at 500 rpm, followed by 10 minutes of incubation with PFA 4% and washed. Slides were stored at 4°C until Ki67 or TUNEL assays were performed.

Briefly, for Ki67 assay slides samples were put in a wet chamber and cells permeabilized with PBT buffer (PBS and 0.2% Triton) for 10 min at room temperature (RT); blocking of non-specific sites was obtained by slide incubation with PBT supplemented with 10% FBS for 30 min at RT. Slides were first incubated overnight at 4°C with the anti-Ki67 antibody (#9129 Cell Signaling, 1:400), washed with PBT and then incubated with Alexa-488 anti-Rabbit secondary antibody (#6441-30 SouthernBiotech, 1:250) for 2h. Slides were washed with PBT and water, mounted with EverBrite Mounting Medium with DAPI (23002, Biotium) and fluorescent microscopic images of proliferative cells (Ki67+) were acquired and analysed with ImageJ software (National Institute of Health, MD, US).

Terminal deoxynucleotidyl transferase (TdT) dUTP Nick-End Labeling (TUNEL) assay was performed using In Situ Cell Death Detection kit, AP (Roche Diagnostics, DE), in order to detect apoptotic cells, according to manufacturer’s protocol. Cells, previously seeded on slides were incubated with a permeabilization solution (0.1% Triton 100X and 0.1% sodium citrate) for 2 min at 4°C, then washed with PBS and incubated with TUNEL mixture (composed by Enzyme Solution added to Label Solution) in a wet chamber for 1h at 37°C. After 3 PBS washes, slides were incubated with Converter AP for 30 min at 37°C and then with Substrate Solution (2% NBT/BCIP Stock solution in NBT/BCIP Buffer) for 10 min at RT and dark. Finally, following PBS washes, mounted with DABCO Mounting Medium and brightfield microscopic images of apoptotic cells (TUNEL+) were acquired and analysed with ImageJ software (National Institute of Health, MD, US).

Both fluorescent and brightfield slides images were acquired by NanoZoomer S60 Digital Slide Scanner (Hamamatsu, Japan) at 20x and 80x magnification and two randomly selected fields for each experimental group at 20x were selected for blinded cells counts by three different operators. Panel images of Ki67+ cells were instead acquired by confocal microscopy A1/A1R (Nikon, Japan) at 60x and 100x magnification. The number of Ki67+ and TUNEL+ cells was normalized on the total cells number per image.

### Subject Recruitment and Sampling for gut microbiota profiling

For this study, 23 RSTS subjects and 16 healthy siblings were enrolled. All subjects were recruited in collaboration with the Italian family RSTS association “Associazione RTS Una Vita Speciale ONLUS”. For both patients and controls, exclusion criteria were: treatments with antibiotic and/or probiotic/prebiotic assumption during the previous 3 months. For RSTS patients, inclusion criteria were: confirmed clinical diagnosis with (20/23) or without (3/23) demonstrated *CREBBP*/*EP300* mutation. RSTS diagnosis of all patients was confirmed by an expert geneticist (DM) and genetic tests were performed in our laboratory (CG).

In conjunction with the stool sample collection, a 3-days dietary survey (preceding the sample collection) was filled by caregivers. Dietary food records were processed using a commercially available software (ePhood V2, Openzone, Bresso, Italy).

The study was approved by Ethics Committee of San Paolo Hospital in Milan (Comitato Etico Milano Area 1, Protocol number 2019/EM/076); written informed consent was obtained from enrolled subjects or caregivers.

### Bacterial DNA Extraction and 16S rRNA Gene Sequencing of human gut microbiota

Bacterial genomic DNA in stool samples was extracted as previously described (Verduci et al., 2018) by using the Spin stool DNA kit (Stratec Molecular, Berlin, Germany), according to the manufacturer’s instructions. Briefly, after homogenizing faecal samples in the lysis buffer for inactivating DNases, Zirconia Beads II were added for a complete lysis of bacterial cells by using TissueLyser LT. Bacterial lysates were then mixed with InviAdsorb reagent, a step designed to remove PCR inhibitors. Bacterial DNA was eventually eluted in 100 μL of buffer. 25 ng of extracted DNA was used to construct the sequencing library. The V3–V4 hypervariable regions of the bacterial 16S rRNA were amplified with a two-step barcoding approach according to the Illumina 16S Metagenomic Sequencing Library Preparation (Illumina, San Diego, CA, USA). Library quantification was determined using the Agilent 2100 Bioanalyzer System (Agilent, Santa Clara, CA, USA); libraries were pooled and sequenced on a MiSeq platform (Illumina) in a 2 × 250 bp paired-end run. Obtained 16S rRNA gene sequences were analysed using PANDAseq (Masella et al., 2012), and low-quality reads filtered and discarded. Reads were, then, processed using the QIIME pipeline (release 1.8.0; Caporaso et al., 2010) and clustered into Operational Taxonomic Unit (OTUs) at 97% identity level and discarding singletons (i.e.: OTUs supported by only 1 read across all samples) as likely chimeras. Taxonomic assignment was performed via the RDP classifier (Wang et al., 2007) against the Greengenes database (version 13_8; ftp://greengenes.microbio.me/greengenes_release/gg_13_8_otus), with a 0.5 identity threshold. Alpha-diversity was computed using the Chao1, number of OTUs, Shannon diversity, and Faith’s Phylogenetic Diversity whole tree (PD whole tree) metrics throughout the QIIME pipeline. Beta-diversity was assessed by weighted and unweighted UniFrac distances (Lozupone et al., 2011) and principal coordinates analysis (PCoA).

### Fecal short chain fatty acid quantification

Concentrations of acetate, propionate, iso-butyrate, butyrate and iso-valerate were assessed according to Bassanini et al. (Bassanini et al., 2019). The measurement of SCFAs was performed by gas chromatography, using a Varian model 3400 CX Gas chromatograph fitted with FID detector, split/splitless injector and a SPB-1 capillary column (30m × 0.32mm ID, 0.25 μm film thickness; Supelco, Bellefonte, PA, USA). Calibration curves of SCFAs in concentration between 0.25 and 10 mM were constructed to obtain SCFAs quantification, and 10 mM 2-ethylbutyric acid was used as internal standard. Results are expressed as mg/g of dry weight of faeces.

### Statistical analysis

Data were analysed using Prism software (GraphPad Software) and expressed as mean ± SD. Student’s t tests were used to compare means between groups in western blot analyses, AlphaLISA, Ki67 and TUNEL assays (*Drosophila* and LCLs acetylation, proliferation and death rate), with p<0.05 considered significant (*p<0.05; **p<0.01; ***p<0.001); correlation between HDACi-induced acetylation and proliferative or apoptotic cells was calculated using Pearson correlation coefficient (−1<r<1) and Pearson correlation p value, significant for p<0.05.

For microbiota analysis, statistical evaluation among alpha-diversity indices was performed by a non-parametric Monte Carlo-based test, using 9999 random permutations. PERMANOVA (adonis function) in the R package vegan (version 2.0-10; Oksanen Jari et al., 2013; last access March 25, 2020) was used to compare the microbial community structure of RSTS and HD subjects. Comparisons of the two groups were performed through the MATLAB software (Natick, MA, USA; version 2008b) using Student’s t-test for normally distributed variables and Wilcoxon test for non-normally distributed variables. For evaluating differences in relative abundances of bacterial groups, a Mann-Whitney U-test was performed. P-values <0.05 were considered as significant for each analysis.

## Data Availability

Sequencing data of 16S rRNA amplicons have been deposited in NCBI Short-Read Archive (SRA) under accession number PRJNA616211 (http://www.ncbi.nlm.nih.gov/bioproject/PRJNA616211)

## ACKNOWLEDGMENTS

We thank the patients’ families for participating in this study. CG thanks the Italian Association of Rubinstein-Taybi patients “RTS Una Vita Speciale ONLUS” for its support. This work belongs to the ERN ITHACA network (DM). Microscopy observations were carried out at The Advanced Microscopy Facility Platform - UNItech NOLIMITS - University of Milan.

## AUTHOR CONTRIBUTIONS

VM, EB, CG conceived the project; EDF, EO, PG and CP performed the experiments; CC and MS performed bioinformatics analyses; EAC and GB contributed to molecular analyses; DM contributed to patient recruitment and clinical assessment; TV provided flies reagents and animals and design of *in vivo* experiments; EDF, EO, VM, EB and CG performed data analysis and interpretation; EDF, EO, VM, EB, CG wrote the manuscript; EV, DM and TV provided guidance in the manuscript revision. All authors: approved the manuscript.

## COMPLIANCE WITH ETHICAL STANDARDS

All the studies involving patients were approved by Ethics Committee of San Paolo Hospital in Milan (Comitato Etico Milano Area 1, Protocol number 2019/EM/076), and in accordance with the 1964 Helsinki declaration and its later amendments or comparable ethical standards. Written informed consent of patients or caregivers were collected for biological samples studies.

## SUPPLEMENTARY MATERIAL

**Supplementary file S1. RSTS LCLs used for *in vitro* treatments**

**Supplementary file S2. Conditions of *in vitro* treatments used on LCLs**

**Supplementary file S3. Insight on single-RSTS LCLs histone acetylation**

**Supplementary file S4. Cell proliferation and cell death rate of RSTS LCLs upon HDAC inhibitors exposure.**

**Supplementary file S5. Correlation between HDACi-induced acetylation versus cell proliferation and apoptosis in RSTS LCLs.**

**Supplementary file S6. Nutritional values of the enrolled patients**

**Supplementary file S7. RSTS gut microbiota analysis**

**Supplementary file S8. Gut microbiota composition in HD and RSTS subjects**

## REFERENCES

Akimaru H, Chen Y, Dai P, Hou D-X, Nonaka M, Smolik SM, Armstrong S, Goodman RH, Ishii S. 1997. Drosophila CBP is a co-activator of cubitus interruptus in hedgehog signalling. Nature 386:735–738. doi:10.1038/386735a0

Alarcón JM, Malleret G, Touzani K, Vronskaya S, Ishii S, Kandel ER, Barco A. 2004. Chromatin Acetylation, Memory, and LTP Are Impaired in CBP+/-Mice. Neuron 42:947–959. doi:10.1016/j.neuron.2004.05.021

Bach Knudsen KE, Lærke HN, Hedemann MS, Nielsen TS, Ingerslev AK, Gundelund Nielsen DS, Theil PK, Purup S, Hald S, Schioldan AG, Marco ML, Gregersen S, Hermansen K. 2018. Impact of Diet-Modulated Butyrate Production on Intestinal Barrier Function and Inflammation. Nutrients 10:1499. doi:10.3390/nu10101499

Bassanini G, Ceccarani C, Borgo F, Severgnini M, Rovelli V, Morace G, Verduci E, Borghi E. 2019. Phenylketonuria Diet Promotes Shifts in Firmicutes Populations. Front Cell Infect Microbiol 9:101. doi:10.3389/fcimb.2019.00101

Benjamin JS, Pilarowski GO, Carosso GA, Zhang L, Huso DL, Goff LA, Vernon HJ, Hansen KD, Bjornsson HT. 2017. A ketogenic diet rescues hippocampal memory defects in a mouse model of Kabuki syndrome. Proc Natl Acad Sci U S A 114:125–130. doi:10.1073/pnas.1611431114

Bercik P, Denou E, Collins J, Jackson W, Lu J, Jury J, Deng Y, Blennerhassett P, MacRi J, McCoy KD, Verdu EF, Collins SM. 2011. The intestinal microbiota affect central levels of brain-derived neurotropic factor and behavior in mice. Gastroenterology 141:599–609, 609–3. doi:10.1053/j.gastro.2011.04.052

Bjornsson HT. 2015. The Mendelian disorders of the epigenetic machinery. Genome Res. doi:10.1101/gr.190629.115

Borgo F, Verduci E, Riva A, Lassandro C, Riva E, Morace G, Borghi E. 2017. Relative Abundance in Bacterial and Fungal Gut Microbes in Obese Children: A Case Control Study. Child Obes 13:78–84. doi:10.1089/chi.2015.0194

Broderick NA, Lemaitre B. 2012. Gut-associated microbes of Drosophila melanogaster. Gut Microbes. doi:10.4161/gmic.19896

Capo F, Wilson A, Di Cara F. 2019. The intestine of Drosophila melanogaster: An emerging versatile model system to study intestinal epithelial homeostasis and host-microbial interactions in humans. Microorganisms. doi:10.3390/microorganisms7090336

Caporaso JG, Kuczynski J, Stombaugh J, Bittinger K, Bushman FD, Costello EK, Fierer N, P?a AG, Goodrich JK, Gordon JI, Huttley GA, Kelley ST, Knights D, Koenig JE, Ley RE, Lozupone CA, McDonald D, Muegge BD, Pirrung M, Reeder J, Sevinsky JR, Turnbaugh PJ, Walters WA, Widmann J, Yatsunenko T, Zaneveld J, Knight R. 2010. QIIME allows analysis of high-throughput community sequencing data. Nat Methods. doi:10.1038/nmeth.f.303

Chan HM, La Thangue NB. 2001. p300/CBP proteins: HATs for transcriptional bridges and scaffolds. Cell Sci 114:2363–73.

Chang MC, Chen YJ, Lian YC, Chang BE, Huang CC, Huang WL, Pan YH, Jeng JH. 2018. Butyrate stimulates histone H3 acetylation, 8-isoprostane production, RANKL expression, and regulated osteoprotegerin expression/secretion in MG-63 osteoblastic cells. Int J Mol Sci 19. doi:10.3390/ijms19124071

Chriett S, Dąbek A, Wojtala M, Vidal H, Balcerczyk A, Pirola L. 2019. Prominent action of butyrate over β-hydroxybutyrate as histone deacetylase inhibitor, transcriptional modulator and anti-inflammatory molecule. Sci Rep 9:742. doi:10.1038/s41598-018-36941-9

Cobos SN, Bennett SA, Torrente MP. 2019. The impact of histone post-translational modifications in neurodegenerative diseases. Biochim Biophys Acta - Mol Basis Dis. doi:10.1016/j.bbadis.2018.10.019

Davie JR. 2003. Inhibition of Histone Deacetylase Activity by Butyrate. J Nutr 133:2485S–2493S. doi:10.1093/jn/133.7.2485s

De Schutter H, Nuyts S. 2009. Radiosensitizing Potential of Epigenetic Anticancer Drugs. Anticancer Agents Med Chem 9:99–108. doi:10.2174/187152009787047707

Douglas AE. 2018. The Drosophila model for microbiome research. Lab Anim (NY) 47:157–164. doi:10.1038/s41684-018-0065-0

Dutto I, Scalera C, Prosperi E. 2018. CREBBP and p300 lysine acetyl transferases in the DNA damage response. Cell Mol Life Sci. doi:10.1007/s00018-017-2717-4

Fast D, Duggal A, Foley E. 2018. Monoassociation with Lactobacillus plantarum Disrupts Intestinal Homeostasis in Adult Drosophila melanogaster. MBio 9. doi:10.1128/mBio.01114-18

Fergelot P, Van Belzen M, Van Gils J, Afenjar A, Armour CM, Arveiler B, Beets L, Burglen L, Busa T, Collet M, Deforges J, de Vries BBA, Dominguez Garrido E, Dorison N, Dupont J, Francannet C, Garciá-Minaúr S, Gabau Vila E, Gebre-Medhin S, Gener Querol B, Geneviève D, Gérard M, Gervasini CG, Goldenberg A, Josifova D, Lachlan K, Maas S, Maranda B, Moilanen JS, Nordgren A, Parent P, Rankin J, Reardon W, Rio M, Roume J, Shaw A, Smigiel R, Sojo A, Solomon B, Stembalska A, Stumpel C, Suarez F, Terhal P, Thomas S, Touraine R, Verloes A, Vincent-Delorme C, Wincent J, Peters DJM, Bartsch O, Larizza L, Lacombe D, Hennekam RC. 2016. Phenotype and genotype in 52 patients with Rubinstein–Taybi syndrome caused by EP300 mutations. Am J Med Genet Part A 170:3069–3082. doi:10.1002/ajmg.a.37940

Freese K, Seitz T, Dietrich P, Lee SML, Thasler WE, Bosserhoff A, Hellerbrand C. 2019. Histone deacetylase expressions in hepatocellular carcinoma and functional effects of histone deacetylase inhibitors on liver cancer cells in vitro. Cancers (Basel) 11. doi:10.3390/cancers11101587

Gottlicher M, Minucci S, Zhu P, Krämer OH, Schimpf A, Giavara S, Sleeman JP, Lo Coco F, Nervi C, Pelicci PG, Heinzel T. 2001. Valproic acid defines a novel class of HDAC inhibitors inducing differentiation of transformed cells. - PubMed - NCBI. EMBO J.

Grunstein M. 1997. Histone acetylation in chromatin structure and transcription. Nature. doi:10.1038/38664

Heerboth S, Lapinska K, Snyder N, Leary M, Rollinson S, Sarkar S. 2014. Use of epigenetic drugs in disease: an overview. Genet Epigenet 6:9–19. doi:10.4137/GEG.S12270

Heijtz RD, Wang S, Anuar F, Qian Y, Björkholm B, Samuelsson A, Hibberd ML, Forssberg H, Pettersson S. 2011. Normal gut microbiota modulates brain development and behavior. Proc Natl Acad Sci U S A 108:3047–3052. doi:10.1073/pnas.1010529108

Hennekam RCM. 2006. Rubinstein–Taybi syndrome. Eur J Hum Genet 14:981–985. doi:10.1038/sj.ejhg.5201594

José-Enériz ES, Gimenez-Camino N, Agirre X, Prosper F. 2019. HDAC inhibitors in acute myeloid leukemia. Cancers (Basel). doi:10.3390/cancers11111794

Kazantsev AG, Thompson LM. 2008. Therapeutic application of histone deacetylase inhibitors for central nervous system disorders. Nat Rev Drug Discov. doi:10.1038/nrd2681

Kung AL, Rebel VI, Bronson RT, Ch’ng LE, Sieff CA, Livingston DM, Yao TP. 2000. Gene dose-dependent control of hematopoiesis and hematologic tumor suppression by CBP. Genes Dev 14:272–7.

Lindefeldt M, Eng A, Darban H, Bjerkner A, Zetterström CK, Allander T, Andersson B, Borenstein E, Dahlin M, Prast-Nielsen S. 2019. The ketogenic diet influences taxonomic and functional composition of the gut microbiota in children with severe epilepsy. npj Biofilms Microbiomes 5. doi:10.1038/s41522-018-0073-2

Lipska K, Gumieniczek A, Filip AA. 2020. Anticonvulsant valproic acid and other short-chain fatty acids as novel anticancer therapeutics: Possibilities and challenges. Acta Pharm 70:291–301. doi:10.2478/acph-2020-0021

Lopez-Atalaya JP, Gervasini C, Mottadelli F, Spena S, Piccione M, Scarano G, Selicorni A, Barco A, Larizza L. 2012. Histone acetylation deficits in lymphoblastoid cell lines from patients with Rubinstein-Taybi syndrome. J Med Genet 49:66–74. doi:10.1136/jmedgenet-2011-100354

Louis P, Flint HJ. 2009. Diversity, metabolism and microbial ecology of butyrate-producing bacteria from the human large intestine. FEMS Microbiol Lett 294:1–8. doi:10.1111/j.1574-6968.2009.01514.x

Lozupone C, Lladser ME, Knights D, Stombaugh J, Knight R. 2011. UniFrac: an effective distance metric for microbial community comparison. ISME J 5:169–72. doi:10.1038/ismej.2010.133

Masella AP, Bartram AK, Truszkowski JM, Brown DG, Neufeld JD. 2012. PANDAseq: paired-end assembler for illumina sequences. BMC Bioinformatics 13:31. doi:10.1186/1471-2105-13-31

Milani D, Manzoni F, Pezzani L, Ajmone P, Gervasini C, Menni F, Esposito S. 2015. Rubinstein-Taybi syndrome: clinical features, genetic basis, diagnosis, and management. Ital J Pediatr 41:4. doi:10.1186/s13052-015-0110-1

Oike Y, Hata A, Mamiya T, Kaname T, Noda Y, Suzuki M, Yasue H, Nabeshima T, Araki K, Yamamura K. 1999. Truncated CBP protein leads to classical Rubinstein-Taybi syndrome phenotypes in mice: implications for a dominant-negative mechanism. Hum Mol Genet 8:387–96. doi:10.1093/hmg/8.3.387

Oksanen Jari, Blanchet F. Guillaume, Friendly Michael, Kindt Roeland, Legendre Pierre, McGlinn Dan, Minchin Peter R, O’Hara R B, Simpson Gavin L, Solymos Peter, Stevens M Henry H, Szoecs Eduard, Wagner Helene. 2013. CRAN - Package vegan. https://cran.r-project.org/web/packages/vegan/index.html

Rubinstein JH, Taybi H. 1963. Broad thumbs and toes and facial abnormalities. A possible mental retardation syndrome. Am J Dis Child 105:588–608.

Ryu J-H, Kim S-H, Lee H-Y, Bai JY, Nam Y-D, Bae J-W, Lee DG, Shin SC, Ha E-M, Lee W-J. 2008. Innate immune homeostasis by the homeobox gene caudal and commensal-gut mutualism in Drosophila. Science 319:777–82. doi:10.1126/science.1149357

Schölz C, Weinert BT, Wagner SA, Beli P, Miyake Y, Qi J, Jensen LJ, Streicher W, McCarthy AR, Westwood NJ, Lain S, Cox J, Matthias P, Mann M, Bradner JE, Choudhary C. 2015. Acetylation site specificities of lysine deacetylase inhibitors in human cells. Nat Biotechnol 33:415–23. doi:10.1038/nbt.3130

Sharon G, Segal D, Ringo JM, Hefetz A, Zilber-Rosenberg I, Rosenberg E. 2010. Commensal bacteria play a role in mating preference of Drosophila melanogaster. Proc Natl Acad Sci U S A 107:20051–6. doi:10.1073/pnas.1009906107

Shin SC, Kim S-H, You H, Kim B, Kim AC, Lee K-A, Yoon J-H, Ryu J-H, Lee W-J. 2011. Drosophila microbiome modulates host developmental and metabolic homeostasis via insulin signaling. Science 334:670–4. doi:10.1126/science.1212782

Simon GM, Cheng J, Gordon JI. 2012. Quantitative assessment of the impact of the gut microbiota on lysine epsilon-acetylation of host proteins using gnotobiotic mice. Proc Natl Acad Sci U S A 109:11133–8. doi:10.1073/pnas.1208669109

Soragni E, Gottesfeld JM. 2016. Translating HDAC inhibitors in Friedreich’s ataxia. Expert Opin orphan drugs 4:961–970. doi:10.1080/21678707.2016.1215910

Spartalis E, Athanasiadis DI, Chrysikos D, Spartalis M, Boutzios G, Schizas D, Garmpis N, Damaskos C, Paschou SA, Ioannidis A, Tsourouflis G, Dimitroulis D, Nikiteas NI. 2019. Histone deacetylase inhibitors and anaplastic thyroid carcinoma. Anticancer Res. doi:10.21873/anticanres.13220

Spena S, Gervasini C, Milani D. 2015. Ultra-Rare Syndromes: The Example of Rubinstein–Taybi Syndrome. J Pediatr Genet 4:177–186. doi:10.1055/s-0035-1564571

Stilling RM, van de Wouw M, Clarke G, Stanton C, Dinan TG, Cryan JF. 2016. The neuropharmacology of butyrate: The bread and butter of the microbiota-gut-brain axis? Neurochem Int. doi:10.1016/j.neuint.2016.06.011

Tanaka Y, Naruse I, Maekawa T, Masuya H, Shiroishi T, Ishii S. 1997. Abnormal skeletal patterning in embryos lacking a single Cbp allele: a partial similarity with Rubinstein-Taybi syndrome. Proc Natl Acad Sci U S A 94:10215–20. doi:10.1073/pnas.94.19.10215

Tarasenko N, Chekroun-Setti H, Nudelman A, Rephaeli A. 2018. Comparison of the anticancer properties of a novel valproic acid prodrug to leading histone deacetylase inhibitors. J Cell Biochem 119:3417–3428. doi:10.1002/jcb.26512

Tarnowski M, Tkacz M, Kopytko P, Bujak J, Piotrowska K, Pawlik A. 2019. Trichostatin A Inhibits Rhabdomyosarcoma Proliferation and Induces Differentiation through MyomiR Reactivation. Folia Biol (Praha) 65:43–52.

Tillhon M, Cazzalini O, Nardo T, Necchi D, Sommatis S, Stivala LA, Scovassi AI, Prosperi E. 2012. p300/CBP acetyl transferases interact with and acetylate the nucleotide excision repair factor XPG. DNA Repair (Amst) 11:844–852. doi:10.1016/j.dnarep.2012.08.001

Uchida S, Shumyatsky GP. 2018. Epigenetic regulation of Fgf1 transcription by CRTC1 and memory enhancement. Brain Res Bull 141:3–12. doi:10.1016/j.brainresbull.2018.02.016

Valor LM, Viosca J, Lopez-Atalaya JP, Barco A. 2013. Lysine acetyltransferases CBP and p300 as therapeutic targets in cognitive and neurodegenerative disorders. Curr Pharm Des 19:5051–64.

Verduci E, Moretti F, Bassanini G, Banderali G, Rovelli V, Casiraghi MC, Morace G, Borgo F, Borghi E. 2018. Phenylketonuric diet negatively impacts on butyrate production. Nutr Metab Cardiovasc Dis 28:385–392. doi:10.1016/j.numecd.2018.01.004

Wang Q, Garrity GM, Tiedje JM, Cole JR. 2007. Naïve Bayesian classifier for rapid assignment of rRNA sequences into the new bacterial taxonomy. Appl Environ Microbiol 73:5261–5267. doi:10.1128/AEM.00062-07

Weinert BT, Narita T, Satpathy S, Srinivasan B, Hansen BK, Schölz C, Hamilton WB, Zucconi BE, Wang WW, Liu WR, Brickman JM, Kesicki EA, Lai A, Bromberg KD, Cole PA, Choudhary C. 2018. Time-Resolved Analysis Reveals Rapid Dynamics and Broad Scope of the CBP/p300 Acetylome. Cell 174:231–244.e12. doi:10.1016/j.cell.2018.04.033

Wong AC-N, Chaston JM, Douglas AE. 2013. The inconstant gut microbiota of Drosophila species revealed by 16S rRNA gene analysis. ISME J 7:1922–32. doi:10.1038/ismej.2013.86

Yao TP, Oh SP, Fuchs M, Zhou ND, Ch’ng LE, Newsome D, Bronson RT, Li E, Livingston DM, Eckner R. 1998. Gene dosage-dependent embryonic development and proliferation defects in mice lacking the transcriptional integrator p300. Cell 93:361–72.

Yuan XG, Huang YR, Yu T, Jiang HW, Xu Y, Zhao XY. 2019. Chidamide, a histone deacetylase inhibitor, induces growth arrest and apoptosis in multiple myeloma cells in a caspase-dependent manner. Oncol Lett 18:411–419. doi:10.3892/ol.2019.10301

