## Supplementary Material for "Exogenous and endogenous HDAC inhibitor effects in Rubinstein-Taybi syndrome models"

[#] Correspondence to:

Cristina Gervasini

Department of Health Sciences

Università degli Studi di Milano

Via Antonio di Rudinì, 8

20142, Milano (IT)

Tel/Fax: +39 02 50323028

ORCID: 0000-0002-1165-7935

**Supplementary file S1.** RSTS LCLs used for epigenetic treatments

| Gene | RSTS LCLs | cDNA change | Protein change | Mutation type | Reference |
|---|---|---|---|---|---|
| ***CREBBP*** | RSTS 114 | c.4485-7G>C | p.(R1428_G1465del)<br>p.(F1379_G1465del) | Splicing | Lopez-Atalaya et al. 2012 |
| | RSTS 120 | c.5837dupC | p.(P1947Tfs*19) | Frameshift | Spena et al. 2015 |
| | RSTS 122 | c.4394+5G>T | p.(R1428_G1465del)<br>p.(F1379_G1465del) | Splicing | Spena et al. 2015 |
| | RSTS 176 | c.4508A>T | p.(Y1503F) | Missense (HAT) | Spena et al. 2015 |
| ***EP300*** | RSTS 25 | c.41_51delinsT | p.(K14Ifs*31) | Frameshift | Negri et al. 2015 |
| | RSTS 39 | c.4640dupA | p.(N1547Kfs*3) | Frameshift | Negri et al. 2016 |
| | RSTS 54 | c.669dupT | p.(Q223Sfs*19) | Frameshift | Negri et al. 2015 |
| | RT010-15 | c.4763T>C | p.(M1588T) | Missense (HAT) | this study |

**Supplementary file S2.** Conditions of epigenetic treatments used on LCLs

| **Treatment** | **TSA** | **SAHA** | **VPA** | **NaB** |
|---|---|---|---|---|
| **Against** | Class I, IIa, IIb HDAC | Class I, IIa, IIb HDAC | Class I (HDAC1, HDAC2, HDAC3) | Class I HDAC |
| **Vehicle** | DMSO | DMSO | $H_2O$ | $H_2O$ |
| **Time** | 2h | 24h | 24h | 24h |
| **Dosage** | 1 - **2** - 5 µM | 1 - **2** - 10 µM | 0,5 - 1 - **2** mM | 1 - 2 - **5** mM |

**Supplementary file S3. Insight on single-RSTS LCLs histone acetylation.** Values of H3K27 acetylation normalized on H3K4 unmodified, explored with AlphaLISA®; on Y-axis values of acetylation upon HDACi are expressed as ratio between the treatment and its vehicle, while on X-axis are listed untreated single LCL or LCLs mean and epigenetic treatments. Graphics show *CREBBP* (RSTS 120, RSTS 122 and RSTS 176) and *EP300* (RSTS 25, RSTS 39, RSTS 54 and RT010-15) LCLs response to HDACi exposure (TSA 2µM, SAHA 2µM, VPA 2mM and NaB 5mM) compared to untreated RSTS means and single-RSTS (in shades of grey). Groups were compared using Student's *t-test* as statistical method (*p<0.05; **p<0.01; ***p<0.001).

### *CREBBP* LCLs acetylation levels

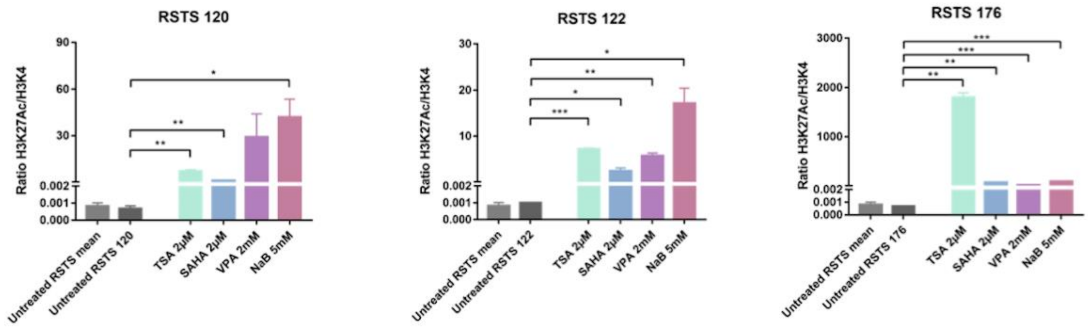


### *EP300* LCLs acetylation levels

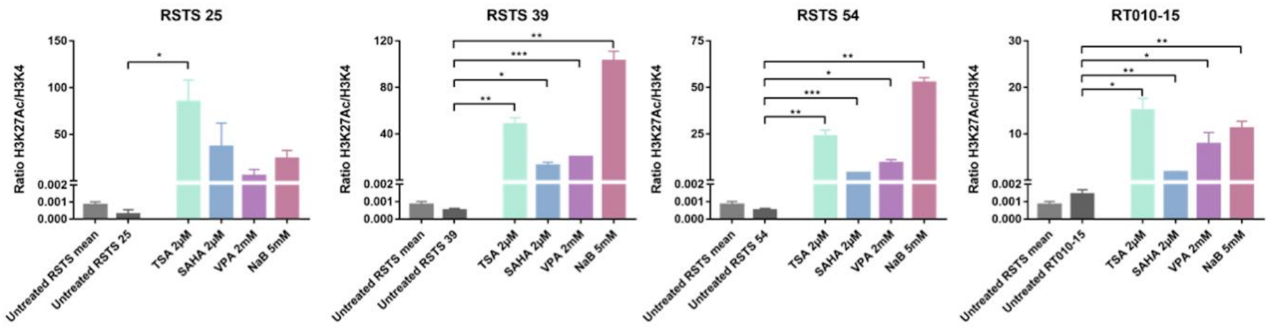

**Supplementary file S4. Cell proliferation and cell death rate of RSTS LCLs upon HDAC inhibitors exposure.** Cell proliferation rate of Ki67 positive cells (% Ki67+ cells, on Y-axis) and cell death rate of TUNEL positive cells (% Apoptotic cells, on Y-axis) of eight RSTS LCLs (*CREBBP* LCLs in shades of red, *EP300* LCLs in shades of pink) after exposure to the four different HDACi (TSA 2μM, SAHA 2μM, VPA 2mM and NaB 5mM) compared to treated HD and RSTS LCLs means (X-axis). Groups were compared using Student's *t-test* as statistical method (*p<0.05; **p<0.01; ***p<0.001).

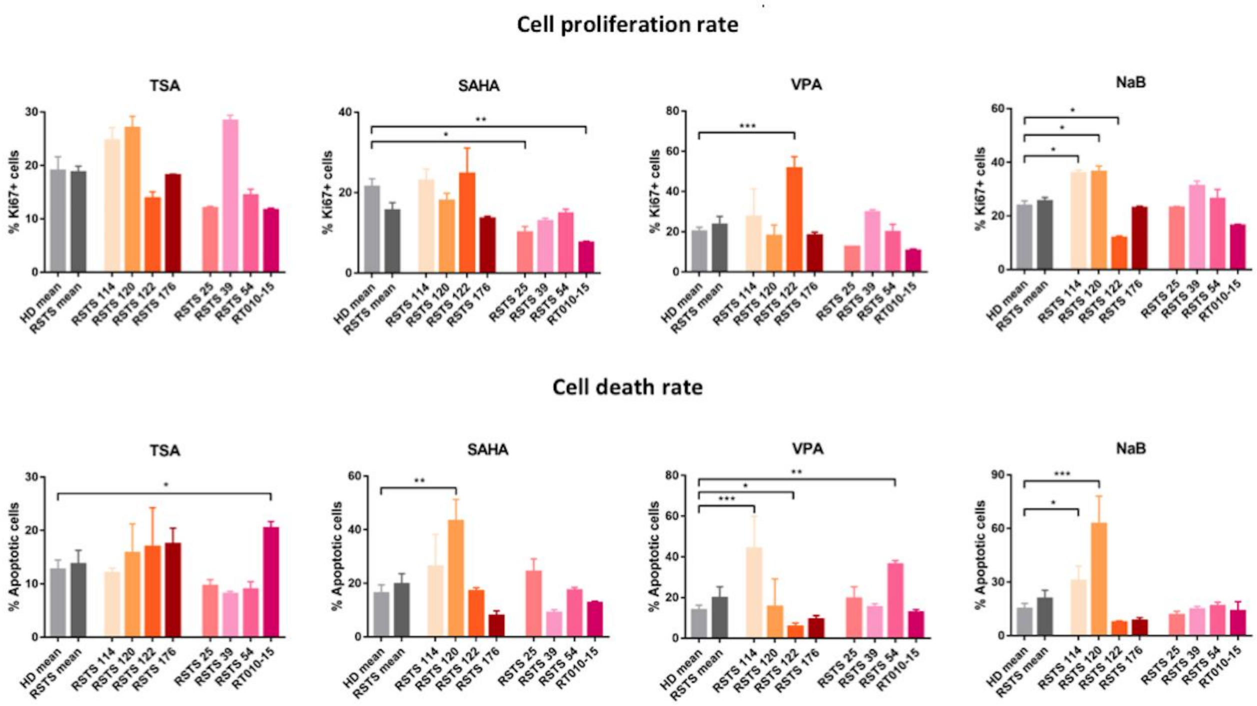

**Supplementary file S5. Correlation between HDACi-induced acetylation versus cell proliferation and apoptosis in RSTS LCLs.** Absence of significant correlation between drug-induced acetylation (X-axis) versus cell proliferation (% Ki67+ cells, on Y-axis) or apoptosis (% Apoptotic cells, on Y-axis) in eight RSTS LCLs (*CREBBP* LCLs in shades of red, *EP300* LCLs in shades of pink) exposed to four different HDACi (TSA 2µM, SAHA 2µM, VPA 2mM and NaB 5mM). Pearson's *r* correlation coefficient was calculated and p<0.05 was considered statistically significant.

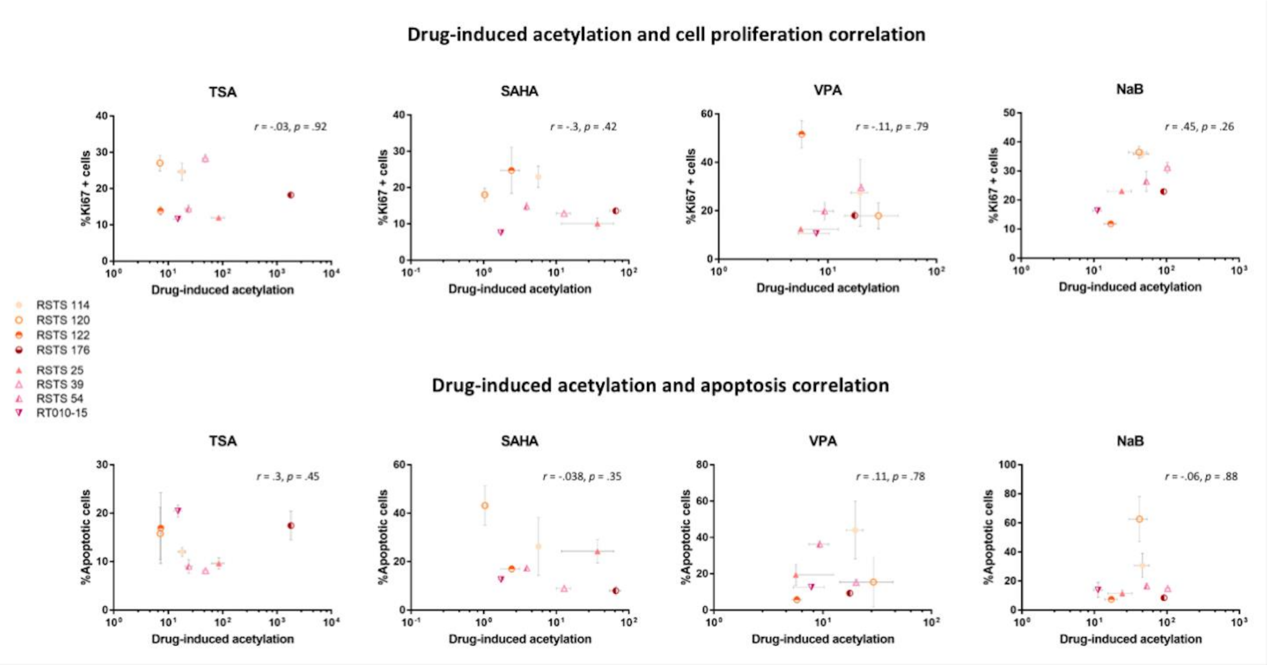

**Supplementary file S6. Nutritional values of the enrolled patients.** Daily dietary intake of energy and macronutrients of in RSTS patients and healthy controls. Values are expressed as mean (standard deviation). p-values <0.05 are considered significant (Mann-Whitney test).

| **Variable** | **HD**<br>**Mean (SD)** | **RSTS**<br>**Mean (SD)** | **p-value** | **Reference values** |
|---|---|---|---|---|
| **Energy intake** | | | | boys:1330-4020 |
| kcal | 1528 (343) | 1185 (294) | 0.0054** | girls:1220-3550<br>kcals (AR) |
| **Proteins** | | | | |
| g | 60.8 (17.97) | 46.22 (13.21) | 0.0079** | 16-50 g (AR) |
| % energy | 15.93 (3.35) | 15.72 (3.29) | 0.8990 | 12-15% (RI) |
| **Lipids** | | | | |
| g | 51.55 (15.05) | 43.73 (13) | 0.0609 | |
| % energy | 30.36 (6.96) | 33.16 (5.38) | 0.1206 | 20-35% (RI) |
| **Carbohydrates** | | | | |
| g | 209.4 (60.81) | 158.8 (41.48) | 0.0054** | |
| % energy | 54.29 (7.62) | 53.65 (5.48) | 0.5626 | 45-60% (RI) |
| **Total fiber** | | | | |
| g | 20.41 (420.05) | 17.33 (13.4) | 0.4369 | |
| g/1000 Kcal | 12.87 (10.35) | 14.54 (9.19) | 0.2065 | 8.40 g/1000 kcal (AI) |

AR. average requirement; RI. reference intake; AI. adequate intake. (Società Italiana di Nutrizione Umana - SINU; “Nutrients and energy reference intake levels for italian population”; 2014, IV revision, Milan, Italy).

**Supplementary file S7. RSTS gut microbiota analysis.** Violin plot showing alpha-diversity calculated with shannon and chao1 metrics. No statistically relevant differences were seen.

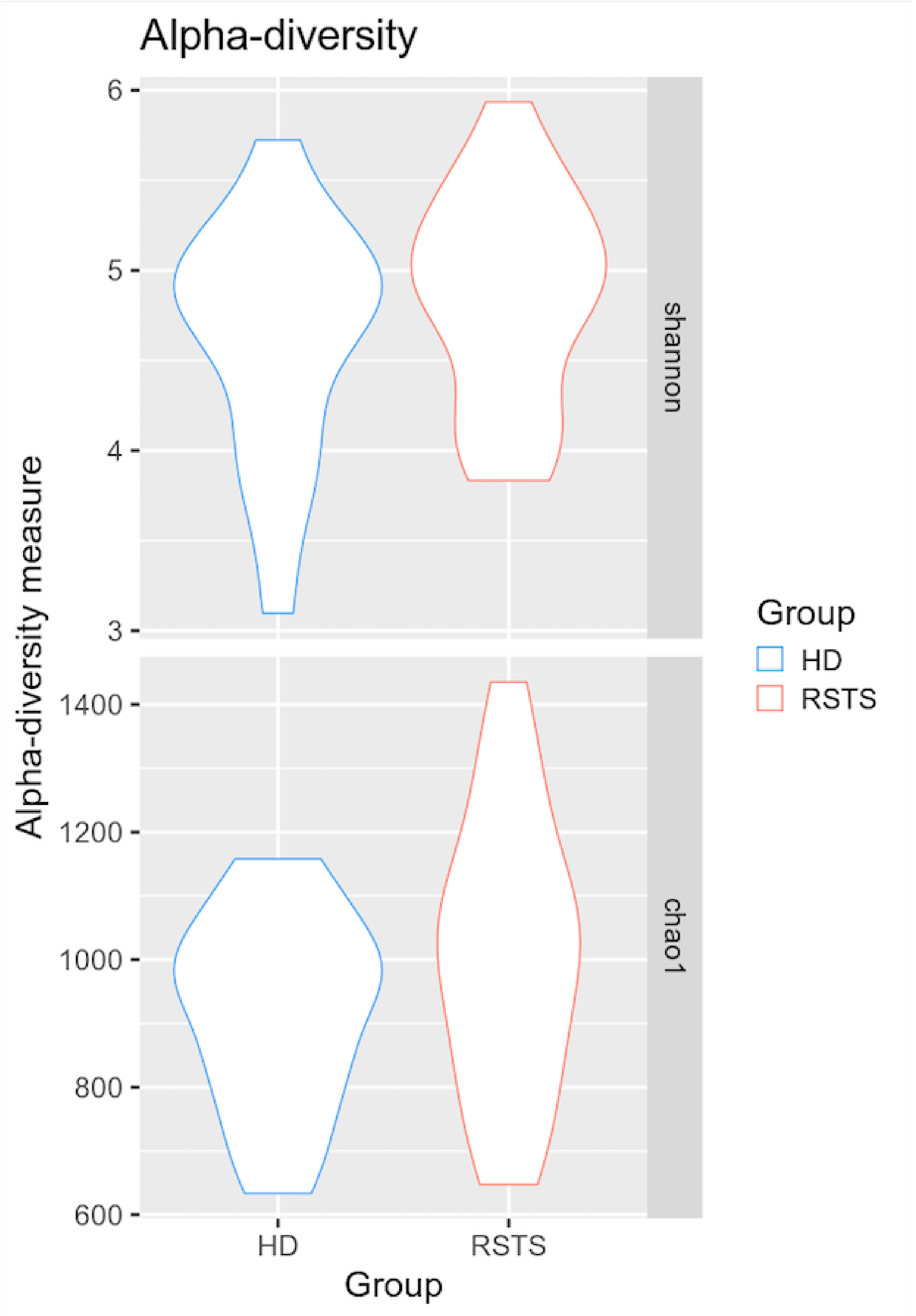

**Supplementary file S8. Gut microbiota composition in HD and RSTS subjects.** Major bacterial groups were organized in three phylogenetic levels (phylum, family, genus) and reported as average relative abundance ± standard deviation. p-values <0.05 were considered significant.

| TAXONOMIC LEVEL | | | HD | RSTS | p-value | |
|---|---|---|---|---|---|---|
| **Phylum** | **Family** | **Genus** | | | | |
| *FIRMICUTES* | | | 73.4 ± 15.6 | 58.5 ± 18.8 | 0.019 | * |
| | *Ruminococcaceae* | | 41.9 ± 15.1 | 32.2 ± 13.9 | 0.049 | * |
| | | *Faecalibacterium* | 9.8 ± 2.2 | 3.3 ± 3.8 | 0.001 | *** |
| | | *Ruminococcus* | 6.4 ± 5.1 | 6.4 ± 4.9 | 0.877 | |
| | | *Oscillospira* | 2.4 ± 2.4 | 5.1 ± 5.0 | 0.007 | ** |
| | | *Ruminococcaceae (other)* | 13.3 ± 15.9 | 8.2 ± 10.2 | 0.746 | |
| | | *Unclass. Ruminococcaceae* | 9.6 ± 9.0 | 9.0 ± 10.1 | 0.525 | |
| | *Lachnospiraceae* | | 16.2 ± 7.2 | 13.1 ± 7.3 | 0.187 | |
| | | *Roseburia* | 5.2 ± 5.8 | 3.4 ± 4.9 | 0.053 | |
| | | *Blautia* | 2.5 ± 3.3 | 1.8 ± 1.3 | 0.855 | |
| | | *Coprococcus* | 2.2 ± 1.4 | 2.0 ± 2.4 | 0.168 | |
| | | *Clostridium* | 1.1 ± 1.6 | 0.6 ± 1.1 | 0.263 | |
| | | *Dorea* | 0.8 ± 0.9 | 0.8 ± 1.0 | 0.855 | |
| | | *Unclass.Lachnospiraceae* | 3.3 ± 3.4 | 2.7 ± 2.1 | 0.471 | |
| | *Veillonellaceae* | | 6.0 ± 6.1 | 5.1 ± 5.4 | 0.703 | |
| | | *Dialister* | 5.1 ± 5.9 | 3.1 ± 4.9 | 0.501 | |
| | *Clostridiaceae* | | 2.4 ± 3.8 | 0.9 ± 1.2 | 0.095 | |
| | | *Clostridium* | 1.1 ± 1.6 | 0.6 ± 1.1 | 0.263 | |
| | *Unclassified Clostridiales* | | 4.8 ± 6.7 | 3.6 ± 5.9 | 0746 | |
| | *Streptococcaceae* | | 1.0 ± 2.0 | 1.8 ± 2.7 | 0.315 | |
| | | *Streptococcus* | 1.0 ± 2.0 | 1.7 ± 2.7 | 0.641 | |
| *BACTEROIDETES* | | | 16.8 ± 14 | 28.7 ± 21 | 0.065 | |
| | *Bacteroidaceae* | | 10.3 ± 10.3 | 21.1 ± 16.3 | 0.021 | * |
| | | *Bacteroides* | 10.3 ± 10.3 | 21.1 ± 16.3 | 0.021 | * |
| | *Rikenellaceae* | | 2.6 ± 2.5 | 3.7 ± 3.3 | 0.220 | |
| | | *Unclass.Rikenellaceae* | 2.5 ± 2.4 | 3.6 ± 3.3 | 0.263 | |
| | *Prevotellaceae* | | 2.0 ± 4.4 | 0.9 ± 3.0 | 0.110 | |

| | | | | | |
|---|---|---|---|---|---|
| | | *Prevotella* | 2.1 ± 4.4 | 0.8 ± 3.0 | 0.115 |
| | *Porphyromonadaceae* | | 0.8 ± 1.4 | 1.6 ± 2.2 | 0.177 |
| | | *Parabacteroides* | 1.4 ± 2.2 | 1.5 ± 2.3 | 0.217 |
| *VERRUCOMICROBIA* | | | 6.8 ± 14.7 | 9.4 ± 10.1 | 0.056 |
| | *Verrucomicrobiaceae* | | 6.8 ± 14.7 | 9.4 ± 10.1 | 0.056 |
| | | *Akkermansia* | 6.8 ± 14.7 | 9.4 ± 10.1 | 0.056 |
| *PROTEOBACTERIA* | | | 1.2 ± 1.5 | 2.1 ± 2.1 | 0.061 |
| | *Enterobacteriaceae* | | 1.0 ± 1.5 | 1.5 ± 2.2 | 0358 |
| | | *Escherichia* | 0.8 ± 1.2 | 1.3 ± 2.2 | 0.263 |
| *ACTINOBACTERIA* | | | 1.6 ± 2.2 | 1.1 ± 1.9 | 0.621 |
| | *Bifidobacteriaceae* | | 1.4 ± 2.2 | 1.0 ± 1.9 | 0.724 |
| | | *Bifidobacterium* | 1.4 ± 2.2 | 1.1 ± 1.9 | 0.724 |
